# Single-step generation of homozygous knock-out/knock-in individuals in an extremotolerant parthenogenetic tardigrade using DIPA-CRISPR

**DOI:** 10.1101/2024.01.10.575120

**Authors:** Koyuki Kondo, Akihiro Tanaka, Takekazu Kunieda

## Abstract

Tardigrades are small aquatic invertebrates known for their remarkable tolerance to diverse extreme stresses. To elucidate the *in vivo* mechanisms underlying this extraordinary resilience, the genetic manipulation methods in tardigrades have long been desired. Despite our prior success in somatic cell gene-editing by microinjecting Cas9 ribonucleoproteins (RNPs) into the body cavity of tardigrades, the generation of gene-edited individuals remained elusive.

In this study, employing an extremotolerant parthenogenetic tardigrade species, *Ramazzottius varieornatus*, we established conditions conductive to generating gene-edited tardigrade individuals. Drawing inspiration from the direct parental CRIPSR (DIPA-CRISPR) technique employed in several insects, we simply injected a concentrated Cas9 RNP solution into the body cavity of parental females shortly before their initial oviposition. This approach yielded gene-edited G0 progeny. Notably, only a single allele was predominantly detected at the target locus for each G0 individual, indicative of homozygous mutations. Through co-injecting single-stranded oligodeoxynucleotides (ssODNs) with Cas9 RNPs, we achieved the generation of homozygously knocked-in G0 progeny and these edited-alleles were inherited by G1/G2 progeny.

This establishment of a simple method for generating homozygous knock-out/knock-in individuals not only facilitates *in vivo* analyses of molecular mechanisms underpinning extreme tolerance but also opens avenues for exploring various topics, including Evo-Devo, in tardigrades.

## Introduction

Tardigrades are microscopic invertebrates living in marine, limnic and limno-terrestrial habitats and all of them require surrounding water to grow and reproduce. There have been more than 1,400 species described so far [1]. Among them, some limno-terrestrial species are known to withstand almost complete loss of water by entering a reversible ametabolic dehydrated state referred to as anhydrobiosis [2]. Dehydrated tardigrades exhibit extraordinary resilience against various extreme stresses including high hydrostatic pressure (7.5 GPa), low temperature (−273 °C), high dose of irradiation and so on [3–7]. The molecular mechanisms underlying their resilient ability has not fully understood. Although some other desiccation-tolerant animals are known to accumulate and utilize non-reducing sugar, trehalose as vitrifying protectants against desiccation [8–10], the accumulation of trehalose is much less or even undetectable in anhydrobiotic tardigrades [11, 12]. Instead, recent studies suggested that tardigrades have and utilize their own unique protective proteins whose expression is abundant and/or significantly induced by desiccation during anhydrobiosis [13–16]. Their functions and roles in the resilient ability have been elucidated largely using heterologous expression and/or *in vitro* systems [14, 15, 17–22]. Although RNAi is feasible and has successfully been used for the elucidation of gene functions in some cases [17, 23, 24], the knock-down efficiency varied depending on the target genes and was not always sufficient. In the previous study, we developed the delivery method of Cas9 ribonucleoproteins (RNPs) to adult tardigrade cells via the microinjection of Cas9 RNPs into the body cavity of tardigrades and the subsequent electroporation and demonstrated that gene editing took place in some somatic cells of the injected tardigrades using a largely transparent tardigrade species, *Hypsibius exemplaris* whose tolerant ability is relatively weak among anhydrobiotic tardigrades [25]. The same study also revealed that the electroporation is not a requisite and the microinjection of Cas9 RNPs into the body cavity is sufficient to induce gene editing in some somatic cells in tardigrades. However, the delivery to germ line cells and the subsequent generation of gene-edited individuals has not yet been achieved. Recently, Shirai *et al*. (2022) developed a new gene editing method termed as direct parental CRISPR (DIPA-CRISPR) in cockroaches and red floor beetles [26]. Using DIPA-CRISPR, the gene-edited progeny (G0) can be obtained by simply injecting Cas9 RNPs into the haemocoel of parental female insects. The injected Cas9 RNPs are assumed to be incorporated to vitellogenic oocytes concomitantly with massive uptake of yolk precursors. In agreement with the assumption, in DIPA-CRISPR it was critical to inject into the females at the appropriate stages during vitellogenesis prior to the first oviposition. Our previous observations that the injection alone was sufficient for the delivery of Cas9 RNPs to induce gene editing in the somatic cells in tardigrades and the successful generation of gene-edited progeny by DIPA-CRISPR in some insects, prompted us to find out the appropriate conditions which enable the generation of the gene-edited tardigrade individuals using DIPA-CRISPR-like method. In this study, we employed an anhydrobiotic and extremotolerant tardigrade *Ramazzottius varieornatus* because its genome sequence is available [15], and lay eggs outside of exuviae, which facilitated us to collect eggs and obtain many individuals at a synchronous age to be injected. We particularly examined two critical parameters, the concentration of Cas9 RNPs and the age of females to be injected, both of which were quite different between our previous somatic cell gene-editing in tardigrades and the original DIPA-CRISPR in insects [25, 26]. By adjusting the conditions, we successfully obtained gene-edited progeny (G0) for two target genes. *R. varieornatus* are parthenogenetic species and lay eggs without mating. We found that most of the obtained gene-edited G0 progenies carried the edited alleles as homozygous. In addition, we found that the simultaneous injection of single-stranded oligodeoxynucleotides (ssODNs) with Cas9 RNPs led to the generation of the knock-in progeny. To our surprise, the gene editing efficiency (GEF) in the knock-in trials was comparable to or even slightly higher than those in the knock-out trials.

This study demonstrated that DIPA-CRISPR like method worked in an extremotolerant parthenogenetic tardigrade, *R. varieornatus*, and the simple injection of Cas9 RNPs (+ knock-in donor if necessary) to parental tardigrades with the appropriate conditions is sufficient to obtain homozygous knock-out/knock-in tardigrade individuals. This gene editing method will substantially promote *in vivo* analysis of molecular mechanisms underlying extreme tolerant ability as well as other research subjects such as Evo-Devo in tardigrades.

## Results

### Determination of Cas9 protein concentration for DIPA-CRISPR in *Ramazzottius varieornatus*

In DIPA-CRISPR, relatively high concentration of Cas9 protein was used as the injection solution (3.3 µg/µL) compared to that of our previous tardigrade study (0.41 µg/µL; Supplementary Table S1) [25], and the lowered concentration of Cas9 protein was reported to decrease the gene editing efficiency [26]. Therefore, we attempted to increase the concentration of Cas9 protein in the injection solution. However, the commercial Cas9 protein solution usually contains relatively high concentration of glycerol (e.g., 50% glycerol in IDT product), which could affect the viability of the injected animals. Accordingly, we first examined how high concentration of glycerol can be tolerated by the injected tardigrades. As shown in Supplementary Table S2, the injection of 20% glycerol solution severely decreased the survival rate to 20%, while the survival rate remained around half (45.5%) when using 15% glycerol solution. We thus chose to use a 15% glycerol concentration, which allows 3.0 µg/µL of Cas9 protein in the injection solution, comparable to those in the original DIPA-CRISPR method [26].

### Generation of gene knock-out tardigrade individuals by DIPA-CRISPR

The experimental scheme of DIPA-CRISPR in tardigrades is shown in Fig. 1. In the original DIPA-CRISPR, the developmental stage of the parents to be injected was one of the most critical parameters for the successful gene editing in progeny [26]. In most cases, the best stage is shortly before the first oviposition, which is consistent with the idea that Cas9 RNPs could be transported to oocytes concomitantly with massive uptake of yolk precursors during vitellogenesis. Given that *R. varieornatus* usually starts to lay eggs around 10 days after hatching [7], we examined the time windows between 5 and 10 days after hatching for the injections to the tardigrades, as younger tardigrades (<5-day-old) seemed too juvenile (immature) and were too small to be injected.

**Fig. 1.**
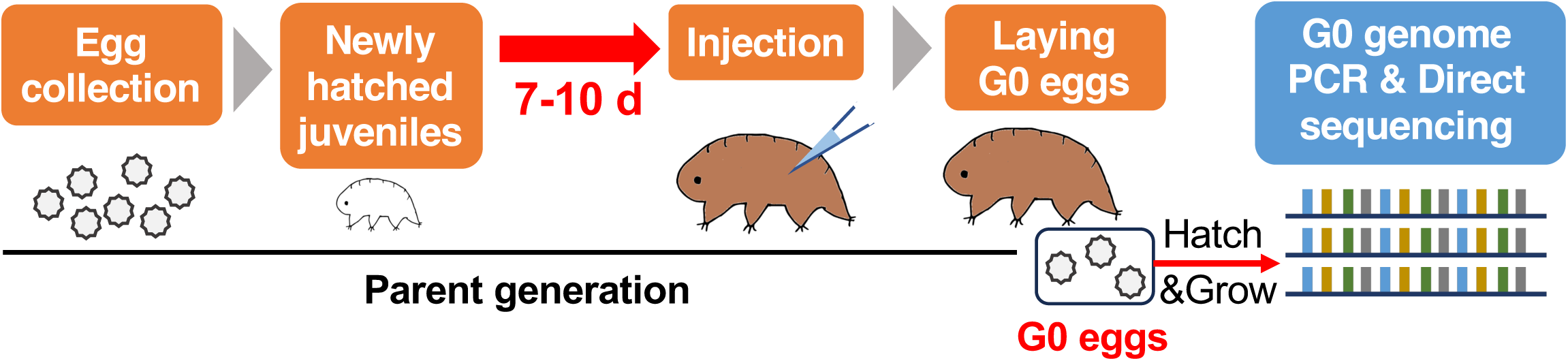
Experimental scheme for DIPA-CRISPR in *R. varieornatus*. The stage of parental females to be injected is a quite important parameter for successful gene-editing in DIPA-CRISPR. To obtain parental females at the defined age to be injected, eggs were collected, and their hatchings were examined daily. Newly hatched juveniles (0-day-old) were separated and reared for the defined period prior to injection. After injection of Cas9 RNPs, the injected tardigrades were reared for 10 days, and the laid eggs (G0 progeny) were collected and reared. Grown G0 individuals were separately subjected to genome DNA extraction and PCR. PCR amplicons were directly analyzed by Sanger sequencing.

We chose the gene RvY_01244 as a target, which encodes an ABC transporter belonging to G subfamily containing the famous *white* gene responsible for a white eye color mutant in *Drosophila melanogaster* [27]. To improve the gene editing efficiency, we synthesized 3 crRNAs (Fig. 2A) and injected RNP solution containing all 3 crRNAs into parental tardigrades of each age from 5- to 10-day-old. We expected some intervening regions among 3 crRNA targets would be deleted from the genome and it would be easily detected by examining the genome PCR amplicon size. In total, we injected 322 parental tardigrades and 102 of them survived for more than 1 day (31.7% survival, Table 1). Using whole bodies of G0 progenies, we successfully obtained genome PCR amplicons about 88 of 184 G0 progenies and found one sample termed *w*-m1 exhibited apparently smaller amplicon size than those expected from the unmodified genome (Fig. 2B). Direct sequencing of the short amplicon revealed the complicated editing in the target locus; i.e., the intervening region (205 bp) between crRNA1 and crRNA2 were lost and the 1,362 bp DNA fragment between crRNA2 and crRNA3 was re-inserted in the inverted orientation (Fig. 2C). It is noteworthy that only short amplicon was obtained from this sample (Fig. 2B) and no mixed peaks were essentially detected in the direct Sanger-sequencing data, suggesting that this tardigrade carried the edited allele as homozygous at the target locus, or carried another mutated allele which suppress the PCR amplification around the target site, e.g., huge deletion. Further direct sequencing of the remaining 87 PCR amplicons identified two additional gene-edited G0 progenies, termed as *w*-m2 and *w*-m3. *w*-m2 carried a 1-nt insertion at the crRNA1 cleavage site (Fig. 2D) and *w*-m3 carried a 3-nt deletion at the crRNA3 cleavage site (Fig. 2E). Again, essentially no mixed peaks were detected in the direct Sanger sequencing data of both samples (Supplementary Fig. S1AB), suggesting both G0 progenies were homozygous at the edited locus. In total, gene editing efficiency (GEF; the proportion of gene-edited individuals out of all sequenced individuals) was 3.4% (Table 1). Overall, the three gene-edited G0 progenies were yielded from the parental animals injected at 8- to 10-day-old. Although the GEF is somewhat lower than the original DIPA-CRISPR, our results indicated that DIPA-CRISPR works and can be used to generate gene knock-out individuals in this tardigrade species.

**Fig. 2.**
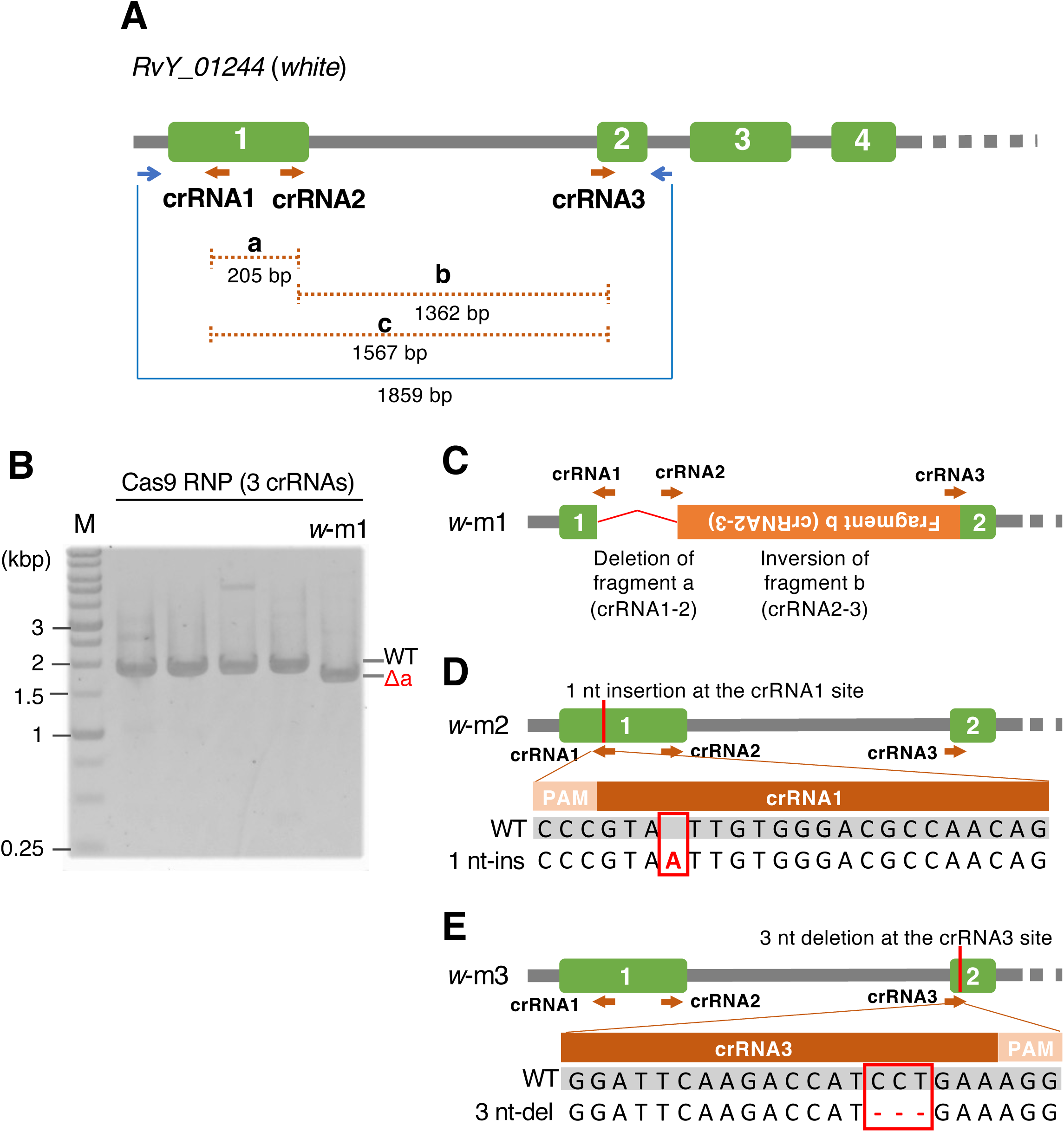
Generation of gene-edited G0 progeny at *white* gene locus. (A) Schematic representation of the gene structure of RvY_01244 (*white*) gene, and the locations of three crRNAs (brown arrows) and genome PCR primers (blue arrows). Green boxes represent exons and gray lines represent introns or intergenic regions. (B) A representative agarose gel image of genome PCR amplicons derived from some G0 progenies. In this gel, the left 4 samples exhibited amplicons at the size expected from unmodified genome (WT) and the right sample termed as *w*-m1 exhibited a single band at shorter size (Δa) than WT, which roughly corresponds the size with the deletion of the fragment a (crNA1-crRNA2). Note: the amplicon at the WT size was not detected in the *w*-m1 sample. (C-E) Gene editing patterns in the three obtained gene-edited G0 individuals, such as a complexed editing (*w*-m1, C), 1-nt insertion (*w*-m2, D) and 3-nt deletion (*w*-m3, E). Each direct Sanger sequencing data of all three gene-edited G0 individuals clearly exhibited a single sequence without mixed peaks (Supplementary Fig. S1). (C) Red bent line represents the deletion of the intervening region between crRNA1 and crRNA2. Orange box represents the intervening DNA fragment between crRNA2 and crRNA3 which was re-inserted in a reverse orientation. (D, E) Schematic representation of the gene-edited location and the comparison of the amplicon sequences with the reference sequence (WT) around the edited sites.

**Table 1.**
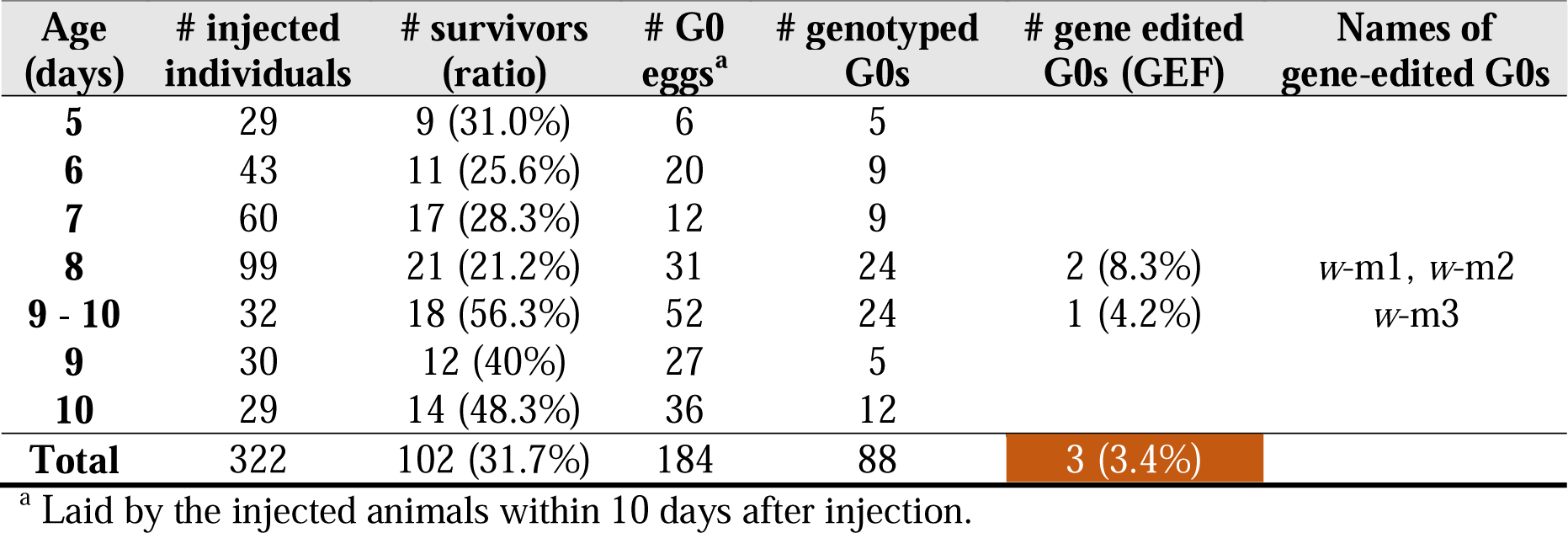
Summary of the gene editing targeting *RvY_01244* (*white*) gene.

| Age (days) | # injected individuals | # survivors (ratio) | # G0 eggs<sup>a</sup> | # genotyped G0s | # gene edited G0s (GEF) | Names of gene-edited G0s |
| --- | --- | --- | --- | --- | --- | --- |
| 5 | 29 | 9 (31.0%) | 6 | 5 |  |  |
| 6 | 43 | 11 (25.6%) | 20 | 9 |  |  |
| 7 | 60 | 17 (28.3%) | 12 | 9 |  |  |
| 8 | 99 | 21 (21.2%) | 31 | 24 | 2 (8.3%) | w-m1, w-m2 |
| 9 - 10 | 32 | 18 (56.3%) | 52 | 24 | 1 (4.2%) | w-m3 |
| 9 | 30 | 12 (40%) | 27 | 5 |  |  |
| 10 | 29 | 14 (48.3%) | 36 | 12 |  |  |
| **Total** | 322 | 102 (31.7%) | 184 | 88 | 3 (3.4%) |  |

<sup>a</sup> Laid by the injected animals within 10 days after injection.

### Editing of trehalose synthesis gene impaired hatchability of next generation

Next, we examined the general applicability of this methodology to other genes. As a next target, we chose *tps-tpp,* a gene responsible for trehalose synthesis. Trehalose is known to play important roles in desiccation tolerance in several anhydrobiotic animals, e.g., nematodes, a sleeping chironomid, and brine shrimps [8–10]. In tardigrades, however, the trehalose production is not a common feature in anhydrobiotic species and the trehalose synthesis gene has been found in only two lineages – superfamily Macrobiotoidea and genus Ramazzottius, both of which were suggested to acquire distinct bacterial trehalose-synthesis genes independently via horizontal gene transfer [28]. *R. varieornatus* has a single *tps-tpp* gene (RvY_13060) which encodes a fusion enzyme of trehalose-6-phosphate synthase (TPS) and trehalose-6-phosphate phosphatase (TPP) and is sufficient to produce trehalose from glucose-6-phosphate and UDP-glucose [15, 28]. We designed 2 crRNAs targeting exon 8 and exon 9 of *tps-tpp* gene respectively, both of which were located within TPS domain (Fig. 3A) and injected RNP solution containing both crRNAs into parental tardigrades of each age from 7- to 10-day-old. As shown in Table 2, we obtained five G0 progeny carrying edited genes. Of those, one individual yielded from the parent injected at 7-day-old and four individuals yielded from those injected at 10-day-old. In total, GEF was 3.4%. In all examined G0 progeny including the gene-edited ones, genome PCR amplicons were essentially detected as a single band in agarose gel electrophoresis (Fig. 3B). Sanger sequencing of the amplicons revealed that four of the five gene-edited G0s carried distinct insertions or deletions without apparent mixed peaks, suggesting these 4 G0 progenies carried homozygous mutation (Fig. 3C, Supplementary Fig. S2A-D). The remaining one exhibited partly mixed peaks in Sanger data which could be interpreted as a mixture of two sequences; the unmodified genome and 1-nt deletion at the cleavage site of crRNA1, though both sequences commonly carried 333bp deletion at the cleavage site of crRNA2 (Fig. 3CD). The peak signals of the unmodified sequence were generally stronger than those of the 1-nt deleted sequence, suggesting that the 1-nt deletion might occur in minor cell population during the development of this G0 progeny resulting in a mosaic organism, while 333bp deletion likely occurred in the oocyte stage resulting as a homozygous mutation.

**Fig. 3.**
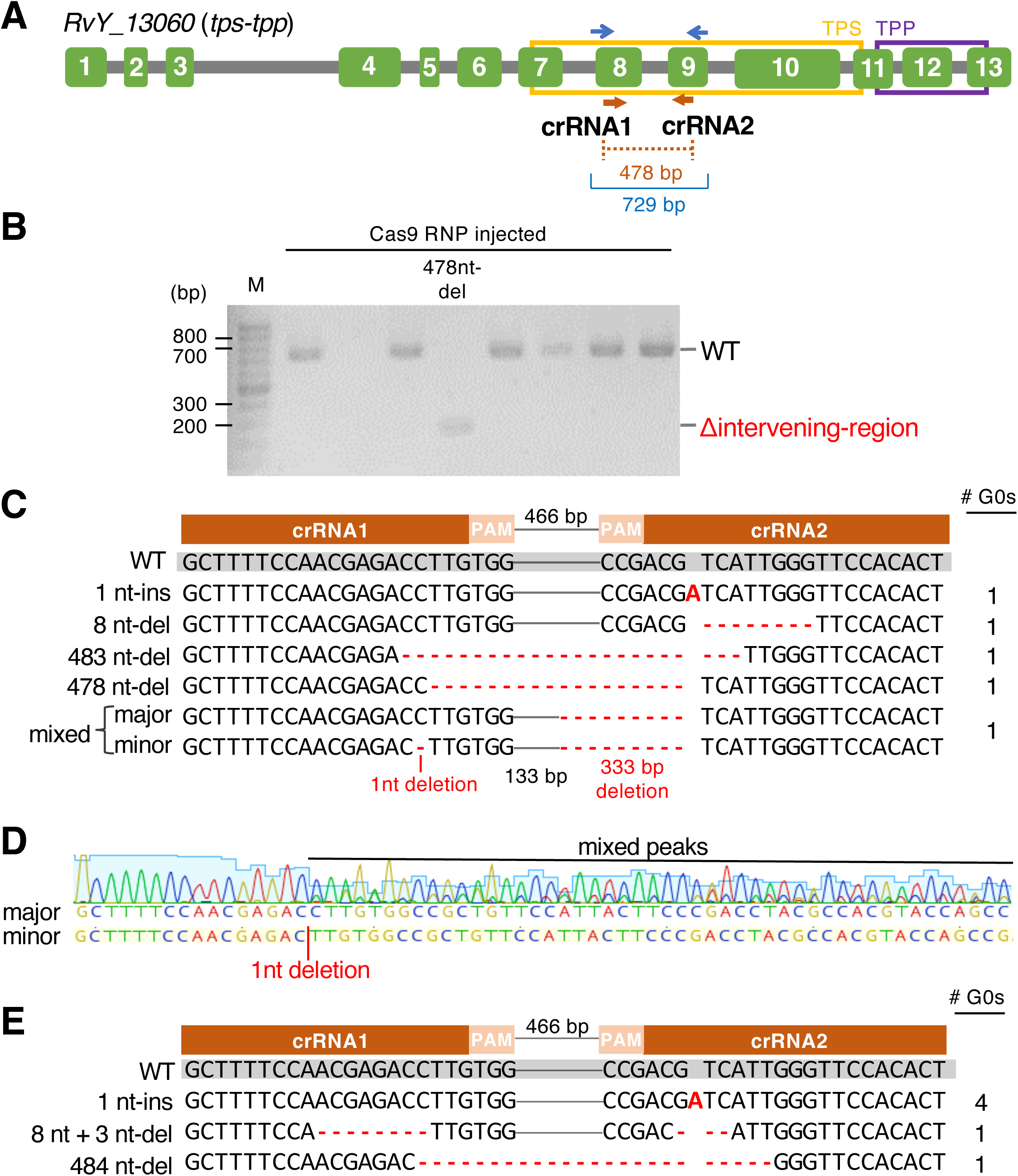
Generation of *tps-tpp* knock-out tardigrades. (A) Schematic representation of gene structure of RvY_13060 (*tps-tpp*), and the locations of two crRNAs (brown arrows) and genome PCR primers (blue arrows). Green boxes represent exons and gray lines represent introns. (B) A representative agarose gel image of genome PCR amplicons from some G0 progenies. WT indicates the amplicon size predicted from unmodified genome and red Δintervening-region indicates the size with the deletion of the intervening region between two crRNAs. The sample labeled as 478nt-del exhibited a single amplicon at the size of 478-nt deletion. (C) Comparison of the amplicon sequences in 5 gene-edited G0 progenies with the reference sequence (WT). The numbers of G0 individuals carrying each editing pattern are shown in the right column. Bold red letters and hyphens indicate insertion and deletions. In Sanger-sequencing data, 4 gene-edited G0 individuals clearly exhibited a single sequence without mixed peaks (Supplementary Fig. S2A-D). The other one exhibited mixed sequences of unmodified one (major) and 1-nt deletion (minor) at the crRNA1 cleavage site, while both of mixed sequences shared the same 333bp deletion around the crRNA2 cleavage site. (D) Electropherograms of direct Sanger sequencing of the gene-edited G0 individual containing mixed peaks. The Sanger data was obtained using the forward primer (left in panel A). There are no mixed peaks in the left portion prior to the putative 1-nt deletion site. In contrast, in the right portion, minor peaks derived from 1-nt deletion sequence were detected with the major peaks corresponding to unmodified sequence. (E) Gene editing patterns in the gene-edited G0 progenies whose G1 eggs were successfully obtained for further analyses. The numbers of G0 individuals carrying each editing pattern are shown in the right. Each G0 individual exhibited only one kind of edited sequence (Supplementary Fig. S2EF), indicative to homozygous mutation. The intervening 466 bp regions between crRNA1 and crRNA2 are shown as thin gray lines.

**Table 2.** Summary of the gene editing targeting *RvY_13060* (*tps-tpp*) gene.

| Age (days) | # injected individuals | # survivors (ratio) | # G0 eggs<sup>a</sup> | # genotyped G0s | # gene edited G0s (GEF) |
| --- | --- | --- | --- | --- | --- |
| 7 | 93 | 32 (34.4%) | 44 | 36 | 1 (2.8%) |
| 8 | 130 | 31 (23.8%) | 54 | 43 |  |
| 9 | 77 | 14 (18.2%) | 17 | 15 |  |
| 10 | 41 | 15 (36.6%) | 73 | 52 | 4 (7.7%) |
| **Total** | 341 | 92 (27.0%) | 188 | 146 | 5 (3.4%) |

<sup>a</sup> Laid by the injected animals within 10 days after injection.

As shown, DIPA-CRISPR worked to generate the *tps-tpp* knock-out mutants in tardigrades. To examine the effects of *tps-tpp* knockout on the tardigrade physiology, we next attempted to establish *tps-tpp* knock-out strains by rearing G0 individual till they lay G1 eggs before sacrificing for genotyping. We again injected Cas9 RNPs with two same crRNAs targeting *tps-tpp* to parental tardigrades of 7- to 10-day-old collectively. After rearing the G0 progenies till they laid G1 eggs, we analyzed the genome sequence of each G0 progeny. As shown in Table 3, we obtained 6 G0 progeny carrying the edited genes among 151 examined individuals (GEF = 4.0%). Of those, four individuals had the same editing, which was 1-nt insertion at the crRNA2 cleavage site (Fig. 3E). The one of two remaining individuals had 8-nt and 3-nt deletion at the crRNA1 and crRNA2 cleavage sites respectively (Fig. 3E) and the other one lost the intervening region between crRNA1 and crRNA2 and neighboring 1 nt + 5 nt (Fig. 3E). Again, in any edited G0 individuals, only a single amplicon was detected in agarose gel electrophoresis, and no mixed peaks were detected in direct Sanger sequencing (Supplementary Fig. S2EF). Each G0 individual carrying the edited genes laid several G1 eggs (2-8 eggs/G0 individual), in total 24 eggs from 6 G0 individuals (Supplementary Table S3). However, unexpectedly, all of G1 eggs from the gene-edited G0s failed to hatch (the hatching rate was 0%; Supplementary Table S3). On the other hand, the hatchability of G1 eggs laid by G0 individuals carrying no-editing was 89.9% (Supplementary Table S4). These observations suggested that the editing of *tps-tpp* gene impaired the hatchability of the G1 progeny in *R. varieornatus*.

**Table 3.** Summary of generation of *tps-tpp* gene-edited G0 individuals laying G1 progeny.

| Age (days) | # G0 eggs<sup>a</sup> | # hatched G0 eggs (ratio) | # genotyped G0s which laid G1 eggs | # gene edited G0s (GEF) |
| --- | --- | --- | --- | --- |
| 7-10 | 182 | 165 (90.7%) | 151 | 6 (4.0%) |

<sup>a</sup> Laid by injected animals within 10 days after injection.

### Establishment of gene knock-in strains by DIPA-CRISPR

CRISPR/Cas9 system including DIPA-CRISPR has been used to generate not only gene knock-out individuals but also knock-in individuals which enables the precise modification of the target genome region as designed. To investigate whether the method above is applicable for gene knock-in in tardigrades, we co-injected single-stranded oligodeoxynucleotides (ssODNs) with Cas9 RNPs into tardigrades of 7- to 10-day-old. We again targeted the gene *white* (RvY_01244). We designed the crRNA near C-terminus of the coding sequence and the ssODN to introduce 11-nt substitutions; 10 of them are synonymous mutations including two mutations in PAM and the other one changes the amino acid from valine (GTG) to methionine (ATG) (Fig. 4A). As shown in Table 4, we obtained five G0 progeny carrying edited genes out of 107 examined G0 individuals (GEF = 4.7%). Three of them exhibited a clear single sequence without mixed peaks in Sanger sequencing, in which every nucleotide at 11 positions was completely substituted as designed in ssODN (Fig. 4B), suggesting they carried the knocked-in allele in a homozygous manner. Sanger sequencing data of another gene-edited individual exhibited a mixture of the fully knocked-in sequence as major and the unmodified (WT) sequence as minor (Fig.4B), suggesting the mosaic nature of the individual. The remainder was in a more complicated situation; it carried 1-nt insertion at the crRNA cleavage site in a homozygous manner, and also exhibited mixed peaks of the knocked-in sequence and the unmodified sequence at two furthest modification sites from the crRNA cleavage site (Fig. 4B).

**Fig. 4.**
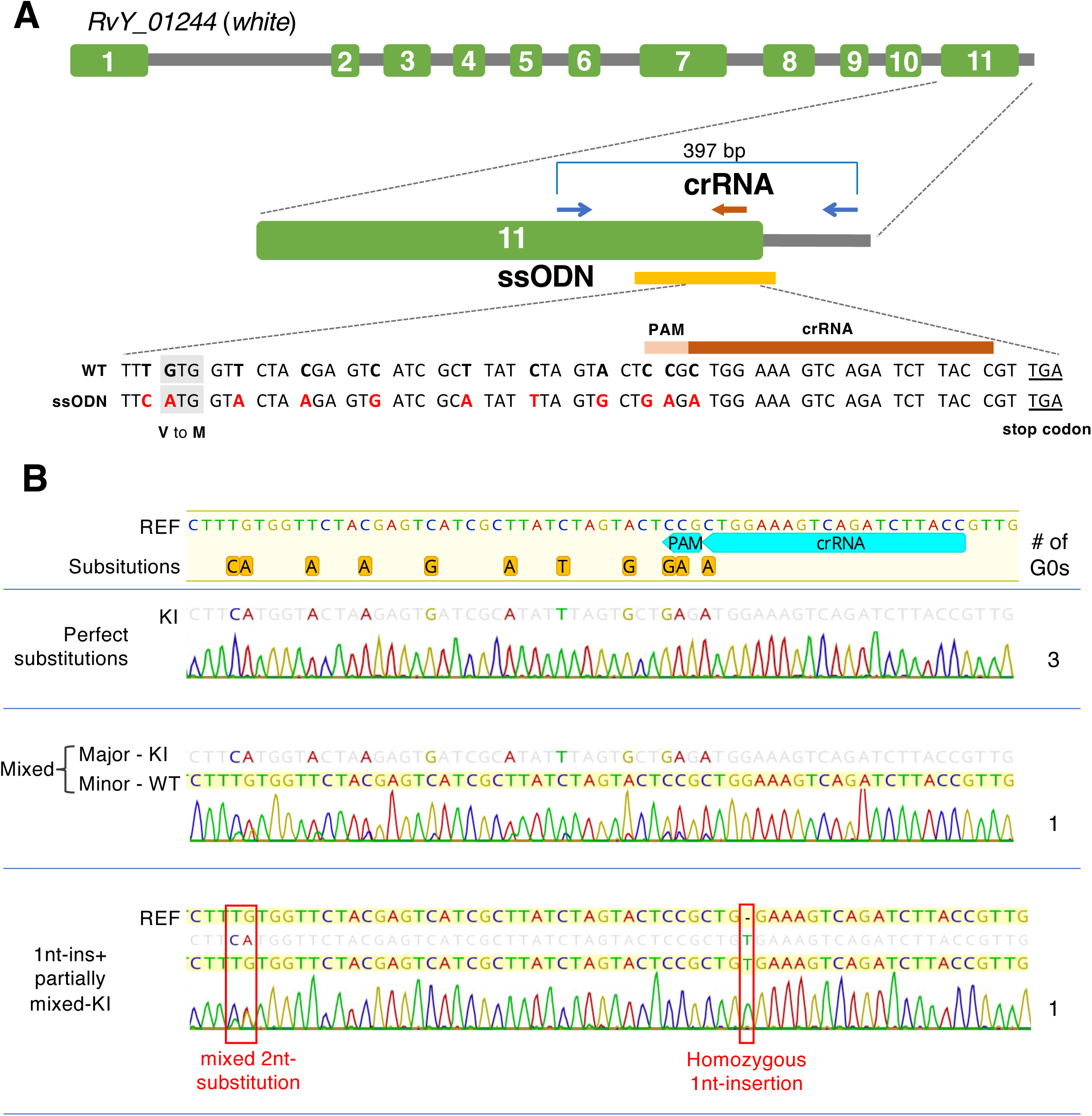
Generation of gene knock-in tardigrades. (A) Schematic representation of gene structure of RvY_01244 (*white*) gene and the locations of crRNA (brown arrows), genome PCR primers (blue arrows) and ssODN (yellow line). Green boxes represent exons and gray lines represent introns and intergenic regions. The sequence of ssODN to introduce the 11 substitutions (red letters) is shown in aligning to reference sequence (WT). (B) Gene editing patterns in the gene-edited G0 progenies obtained by co-injecting ssODN and their representative electropherograms in direct Sanger sequencing of the amplicons. The number of G0 individuals with each editing pattern is shown in the right. The three G0 individuals exhibited a clear single sequence carrying 11-nt substitutions as designed in ssODN (perfect substitutions). One of the other G0 individuals (shown in middle of panel B) exhibited mixed sequences of 11-nt substituted (knocked-in, KI) sequence as major and an unmodified sequence (WT) as minor. The other one (shown in bottom of panel B) carried 1-nt insertion at the cleavage site and also exhibited two consecutive mixed peaks of unmodified and knocked-in sequences at the furthest knock-in position from the cleavage site.

**Table 4.** Summary of knock-in experiments targeting *RvY_01244* (*white* gene).

| Age (days) | # injected individuals | # survivors (ratio) | # G0 eggs<sup>a</sup> | # hatched eggs (ratio) | # genotyped G0s | # gene edited G0 (GEF) |
| --- | --- | --- | --- | --- | --- | --- |
| 7-10 | 210 | 73 (34.8%) | 137 | 125 (91.2%) | 107 | 5 (4.7%) |

<sup>a</sup> Laid by injected animals within 10 days after injection.

To examine whether these edited alleles are heritable, we examined the genotypes of G1/G2 progeny of these gene-edited G0 individuals after propagation. From perfectly knocked-in G0 individuals, one knocked-in strain was successfully propagated and established, in which G2 progeny were confirmed to carry the perfectly knocked-in allele as homozygous (Supplementary Fig. S3). From the G0 individual carrying mixture of knocked-in sequence as major and unmodified sequence as minor, 5 G1 progenies were obtained. Although one G1 progeny could not be genotyped due to amplification failure, the other 4 G1 progenies were confirmed to carry the fully knocked-in sequence as homozygous leading to the establishment of the knocked-in strains in which G2 progeny were confirmed to carry the same knock-in sequence. And thus, the observed knocked-in sequence was successfully inherited by their progeny. From the remaining G0 progeny which carried homozygous 1-nt insertion and mosaic 2-nt knocked-in sequence, one G1 egg was obtained which carried only 1-nt insertion. The detected 2-nt knock-in sequence as mosaic seemed not heritable, while the homozygous 1nt-insertion was heritable.

## Discussion

In this study, we demonstrated that DIPA-CRISPR successfully worked in the extremotolerant parthenogenetic tardigrade. By using this method, homozygous gene knock-out/knock-in tardigrade individuals were successfully generated in a single step. In the original DIPA-CRISPR, the injected Cas9 RNPs are assumed to be incorporated into vitellogenic oocytes concomitantly with the massive uptake of vitellogenins by receptor-mediated endocytosis, thus injecting the individuals at appropriate developmental stages was one of the critical parameters for the successful yielding of gene-edited progeny [26]. As we have no knowledge about vitellogenic process in *R. varieornatus*, we tried to inject parental tardigrades at 5- to 10-day-old which corresponds just before the first oviposition and obtained gene-edited progeny from the parents injected at 7- to 10-day-old. In a related tardigrade species, *H. exemplaris* which belongs to the same taxonomic family of *R. varieornatus*, vitellogenesis proceeds in three distinct modes lasting four days: the first part of the yolk is synthesized by the oocyte itself (autosynthesis); the second part is synthesized by trophocytes and transported to the oocyte through cytoplasmic bridges; and the third part is synthesized outside the ovary (in storage cells) and transported to the oocyte by endocytosis [29]. In this kind of vitellogenesis, the injected Cas9 RNPs could be incorporated into oocytes during the third part of vitellogenesis. In *R. varieornatus*, the germ cells could uptake the injected Cas9 RNPs likewise.

*R. varieornatus* is a diploid parthenogenetic species [7, 15], though the cytological processes of progeny production and the mode of inheritance of genetic materials remained unclear. Ammermann D. (1967) reported the cytological processes of a diploid parthenogenetic reproduction in the related tardigrade species, *H. dujardini* which is a species complex containing the recently redescribed *H. exemplaris* and belongs to the same taxonomic family of *R. varieornatus* [30, 31]. During oogenesis in *H. dujardini*, the female germ cells undergo the first meiosis and daughter cells received the mostly homozygous dyads derived from the meiotic bivalent chromosomes. After that, this dyad disintegrates and the diploidy is recovered in the daughter cells. The meiosis is completed by the following mitosis-like process of the second meiosis which keeps the diploidy. According to this cytological process, the chromosomes of the oocytes are predicted to be largely homozygous except the small possible heterozygous regions which could be derived from chromosomal crossover during the first meiosis. In our genotyping analyses of the edited G0 individuals of *R. varieornatus*, only a single sequence was detected in most direct Sanger sequencing, suggesting that most G0 progeny carried the edited allele in a homozygous manner (Fig. 2B-E, Fig. 3B-E, Fig. 4B; Supplementary Fig. S1, Supplementary Fig. S2). Especially, a similar result was obtained in the case of a very complicated editing of *white* gene by 3 crRNAs, in which one intervening region was deleted and the other intervening fragment was re-inserted in the inverted orientation (Fig. 2C). It is unlikely that Cas9 RNPs independently performed the same complicated editing on both alleles in a germ cell. And thus, this result is very difficult to explain if *R. varieornatus* undergo a clonal (ameiotic) propagation. If the cytological process of parthenogenetic reproduction in *R. varieornatus* is similar to that in *H. dujardini*, it was assumed that CRISPR-Cas9 system would edit the single allele in a germ cell before meiosis and the mutation would be then replicated and transferred to the mature egg cell as homozygous during a meiotic process (Supplementary Fig. S4). This is a good news for researchers, because a homozygous mutant could be obtained in a single step, which significantly facilitate the downstream analyses. In a few cases of our genotyping data, weak mixed peaks were detected (Fig. 3CD, Fig. 4B). We assumed that these minor peaks were likely derived from mosaic mutations which might occur by the delayed action of the remaining Cas9 RNPs in a small cell population during the development of G0 progeny. In general, gene knock-in mediated by homology-directed repair (HDR) tends to occur at much lower rate compared to gene knock-out mediated by nonhomologous end-joining (NHEJ) [32–35], although germ cells are more prone to HDR than somatic cells [36]. In the original DIPA-CRISPR, GEF (the proportion of edited individuals out of the total number of individuals hatched) in knock-in trials was 1.2%, while it was 50.8 - 71.4% in knock-out trials in red flour beetles at the optimized stages [26]. In this study, however, knock-in efficiency was even slightly higher than those of knock-out (4.7% and 3.4 - 4.0%, respectively) and we rarely observed the short indels by NHEJ-mediated repair in the knock-in experiments, suggesting that HDR might be a dominant repair mode in the germ cells of this tardigrade species. It is noteworthy that using *R. variornatus* (in this study), we did not observe the tendency of no-indel NHEJ which was observed in somatic cells of *H. exemplaris* in our previous report [25]. This could be consistent with the relatively low efficiency in NHEJ-dependent repair (knock-out) in this study.

In all *tps-tpp* edited mutants obtained in this study, a frame shift was introduced at the putative cleavage sites of crRNA1 or crRNA2 both of which were located within the TPS domain (Fig. 3A). And thus, the mutated *tps-tpp* gene products likely lost the function of C-terminal region of TPS and whole TPP (Fig. 3A). Because the C-terminal region of TPS is responsible for the binding to the substrate UDP-glucose [37], the TPS activity was likely lost in the edited tardigrades as well. In all *tps-tpp* edited mutants, G0 individuals were able to hatch, grow and lay eggs normally, but no G1 eggs hatched (Supplementary Table S3). The hatchability of G1 eggs was significantly lower in *tps-tpp* edited mutants than in those harboring no-editing in *tps-tpp* gene (*p* = 2.57e-15, Fisher’s exact test). These results suggested that the mutations in *tps-tpp* gene had a maternal effect on the hatchability of the embryos in this tardigrade species. For instance, trehalose could be synthesized in maternal tissue and transported to oocyte and might play important roles in embryogenesis of the progeny, e.g., as an energy reserve. In cockroaches, *Periplaneta americana*, the treatment with trehalase inhibitor validoxylamine A (VAA) inhibited normal oocyte development, indicating that trehalose is necessary for successful oocyte development in this insect species [38]. However, we would like to leave a question open whether trehalose itself plays an important role in tardigrade physiology. Because the TPS-TPP protein of *R. varieornatus* contains extraordinarily long N-terminal region which exhibits sequence similarity with trans-1,2-dihydrobenzene-1,2-diol dehydrogenase. This N-terminal region could be translated even in the *tps-tpp* mutants and the truncated gene products might be harmful and responsible for the observed phenotype in this study, instead of trehalose reduction.

In summary, we have successfully established a method for generating both gene knock-out and knock-in individuals in the anhydrobiotic and extremotolerant tardigrade species, *R. varieornatus*, through adjusting conditions of DIPA-CRISPR. Our findings indicated that the optimal injection window is between 7 and 10 days after hatching aligning with the period shortly before the first oviposition in this species. The simple injection of Cas9 RNPs (with knock-in donor when necessary) to parental tardigrades at the appropriate age is sufficient to obtain the edited progeny. Notably, these progenies predominantly carried the edited allele as homozygous, likely attributed to the meiotic parthenogenetic mode of reproduction. This feature significantly facilitates loss of function analyses downstream. While DIPA-CRISPR was initially developed for insects [26], our study shows its effectiveness in a non-insect, even non-arthropod organism, underscoring the broad applicability of this method to various invertebrate species, including other tardigrades. This method will facilitate *in vivo* analysis of various topics in tardigrades, including molecular mechanisms underlying their renowned extreme tolerance as well as many Evo-Devo subjects.

## Materials and methods

### Tardigrades

*Ramazzottius varieornatus* YOKOZUNA-1 strain was reared as described previously [21]. Briefly, tardigrades were maintained in 1.2% agar plates overlaid with sterilized pure water (Elix Advantage 3 UV, Millipore) containing live chlorella suspension (Recentec) as food at 22 °C. Water and food were replaced once per week. To prepare the staged tardigrades for injection, eggs were collected from culture dishes and were transferred to a new agar dish with food after the cleaning treatment with 1% commercial chlorine bleach. Next day, unhatched eggs and eggshells were removed from the dish to leave only newly hatched juveniles which were labeled as 0-day-old. These juveniles were reared as described above till the appropriate age.

### Preparation of Cas9 complex (RNPs)

S.p. Cas9 Nuclease V3, CRISPR-Cas9 tracrRNA and crRNAs were purchased from IDT. For each target genome region, the list of possible crRNAs and off-target information in *R. varieornatus* were retrieved using CRISPR direct (https://crispr.dbcls.jp/) and on-target scores of crRNAs were also obtained from manufacturer’s web site (https://sg.idtdna.com/site/order/designtool/index/CRISPR_CUSTOM). Based on these information, appropriate crRNAs were selected as described in Supplementary Table S5. Cas9 complex (RNPs) was prepared as essentially described previously with increasing the concentration of each component [25]. Briefly, the mixture of crRNAs and tracrRNA (final 100 µM each) were heated at 95 °C for 5 min, and then gradually cooled into room temperature. In 10 µL scale, 1 µL of PBS (137 mM NaCl, 2.7 mM KCl, 10 mM Na_2_HPO_4_, 1.8 mM KH_2_PO_4_), 3 µL of 10 µg/µL Cas9 protein (IDT, final 3.0 µg/µL), 6 µL of 100 µM gRNA (final 60 µM) were mixed and incubated at room temperature for 15-20 min to assemble Cas9 RNPs. Cas9 RNP solution was kept at −80 °C until use.

### Injections

The injection was performed as described in the previous study [25]. Briefly, the staged animals were anesthetized in 25 mM levamisole (Sigma) and mounted on an injection slide. The injection slide was placed on an inverted differential interference contrast microscope (Axiovert 405 M, Zeiss). Glass capillary needle was prepared from a glass capillary (GD-1, Narishige) using a needle puller (PC-10, Narishige). Cas9 RNP solution was filled into a glass capillary needle and injected into a body cavity of a tardigrade, by using FemtoJet 4i (Eppendorf) with the settings as pi=1500 hPa, ti=0.20 s, pc=50 hPa. Successful injection was confirmed by swelling of the specimen. Injected individuals were recovered onto agar plates with sterilized water and maintained with food until laying eggs (G0). We collected G0 eggs laid within 10 days after injection. Successfully hatched G0 progeny were subjected to genotyping before or after laying eggs (G1).

### Genotyping

Genome PCR was performed as described in the previous study with some modifications [25]. Briefly, genomic DNA was extracted from each G0 individual in 10 µL of protease K solution (500 µg/mL, in 0.5 x KOD FX Neo PCR buffer (TOYOBO)) by incubating at 60 °C for 120 min. Protease K was then inactivated by heating at 95 °C for 15 min. Genome samples were stored at −20 °C until PCR reaction. Primers were designed to include all target sites of crRNAs (Supplementary Table S5). The target regions were amplified from about 0.5 to 2 µL genome samples with using KOD FX Neo (TOYOBO) in 10 µL reaction. PCR amplicons were examined by agarose gel electrophoresis (AGE). Genome PCR products which were confirmed as a single band in AGE were subjected to direct Sanger sequencing after decomposing remaining nucleotides and primers by the treatment with 2U Exonuclease I (NEB) and 0.1U Shrimp Alkaline Phosphatase (NEB).

### Knock-in experiments

ssODN were designed to introduce 11-nt substitutions, two of which mutate PAM sequence (Supplementary Table S5). The designed ssODN (189 bases) were synthesized and purchased from IDT. RNP solution was prepared as described above and 10 µL of RNP solution was mixed with 1 µL of 14.5 µg/µL ssODNs (final 1.3 µg/µL) and used for injection. The solution was kept at −80 °C until use.

## Acknowledgements

This work was supported by JSPS KAKENHI Grant Numbers JP20H04332, JP20K20580, and JP21H05279 to T.K.. A.T. received a Grant-in-Aid for JSPS Fellows (No. 21J11385).

## Author contributions

K.K. and T.K. conceived and designed the study. K.K. performed experiments and analyzed data. A.T. provided materials. K.K. and T.K. wrote the paper and all authors approved the manuscript.

## Data availability

All data have been included in the manuscript and the supplementary information.

