## Supplemental Materials for "Single-step generation of homozygous knock-out/knock-in individuals in an extremotolerant parthenogenetic tardigrade using DIPA-CRISPR"

**A** *w*-m2

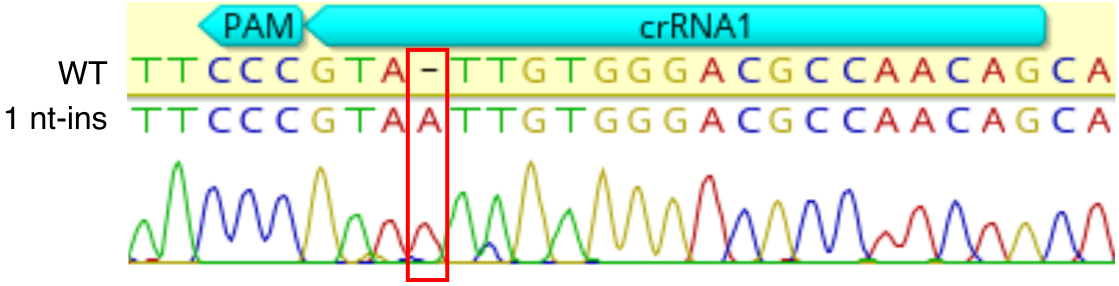


**B** *w*-m3

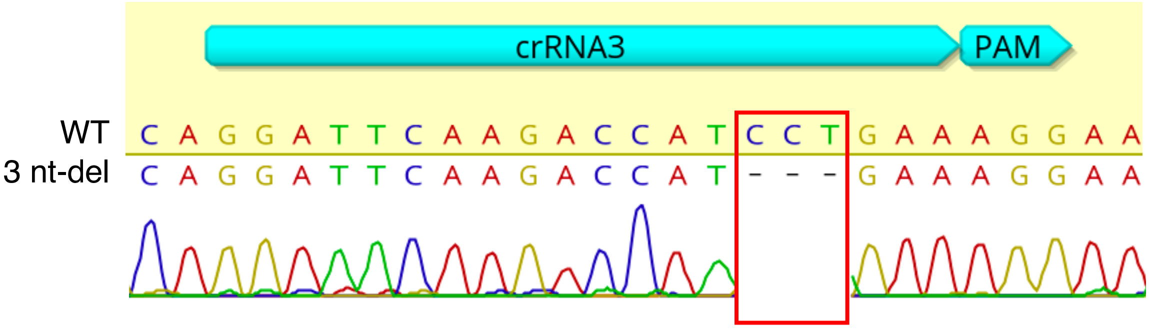


**Supplementary Fig. S1**

Electropherograms in direct Sanger sequencing of genome PCR amplicons obtained from G0 individuals carrying gene-editing in RvY_01244 (*white*) gene locus. (A) *w*-m2 carrying 1-nt insertion. (B) *w*-m3 carrying 3-nt deletion. All sequencing data exhibited clearly distinct peaks without mixture, suggesting that each G0 individual carrying the mutated allele in a homozygous manner.

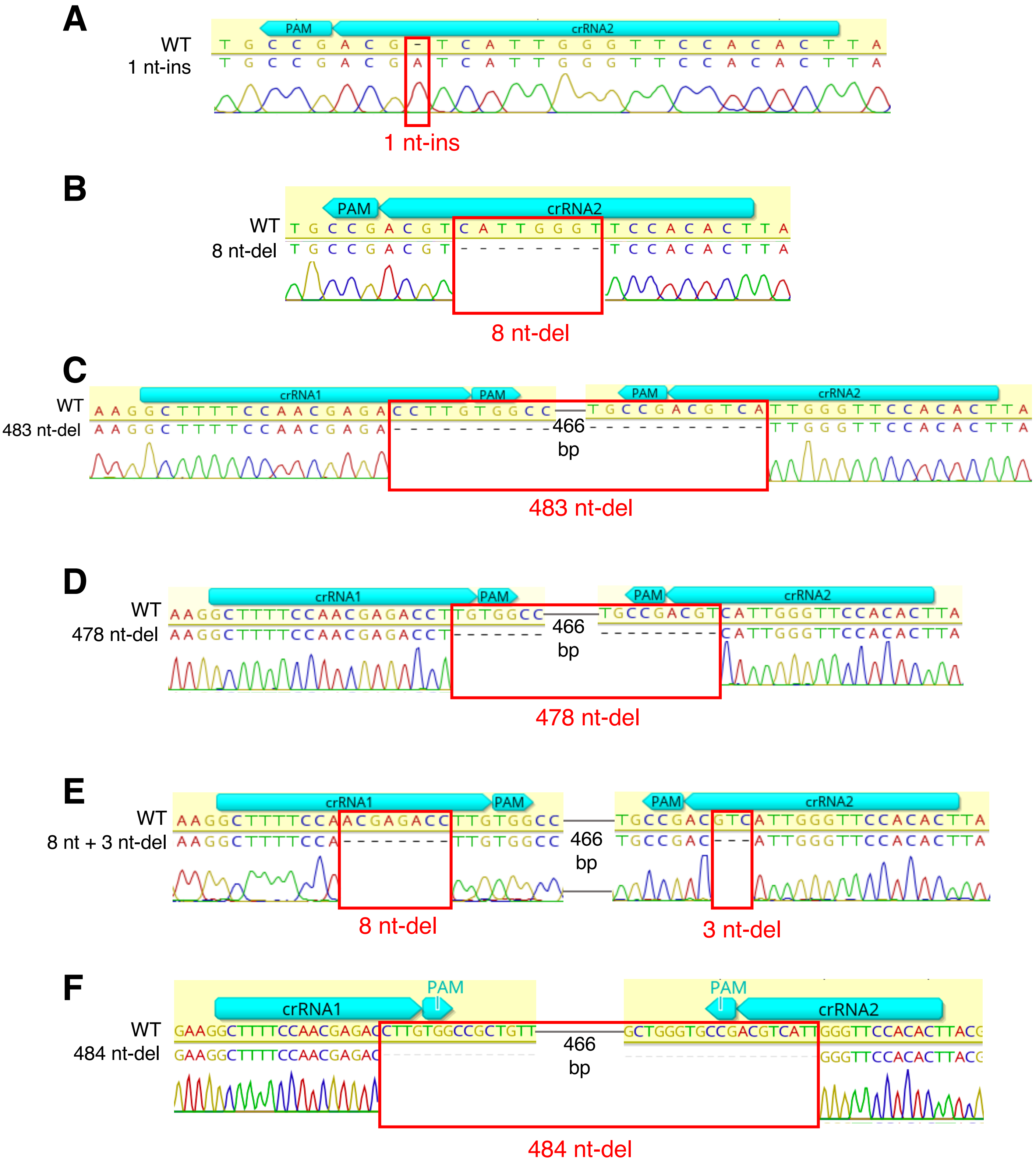


**Supplementary Fig. S2**

Electropherograms in direct Sanger sequencing of genome PCR amplicons obtained from G0 individuals carrying gene-editing in RvY_13060 (*tps-tpp*) gene locus.
Data corresponds to Figure 3C (A, 1 nt-ins; B, 8 nt-del; C, 483 nt-del; D, 478-del) and Figure 3E (E, 8 nt + 3 nt-del; F, 484 nt-del). No mixed peaks were detected, suggesting all G0 individuals are homozygous mutants.

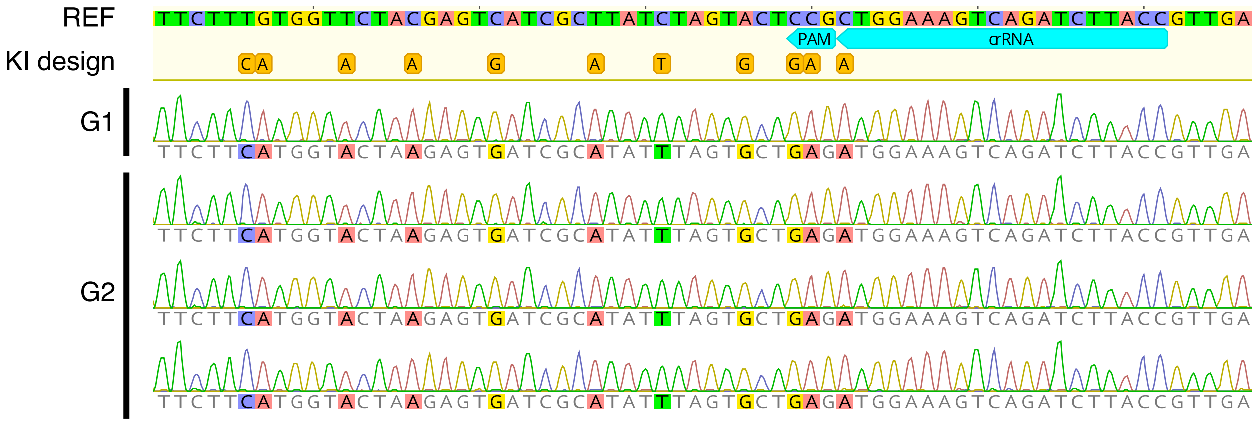


### Supplementary Fig. S3

Representative electropherograms exhibiting perfect substitutions in a G1 individual and three G2 individuals derived from a perfectly knocked-in G0 individual, indicating the heritability of the knock-in sequence. REF represents the sequence of the unmodified genome, and the designed substitutions in ssODN are shown as KI design. The substituted bases are shown as highlighted. All progeny exhibited clear single sequences without mixed peaks, indicative of homozygous mutation.

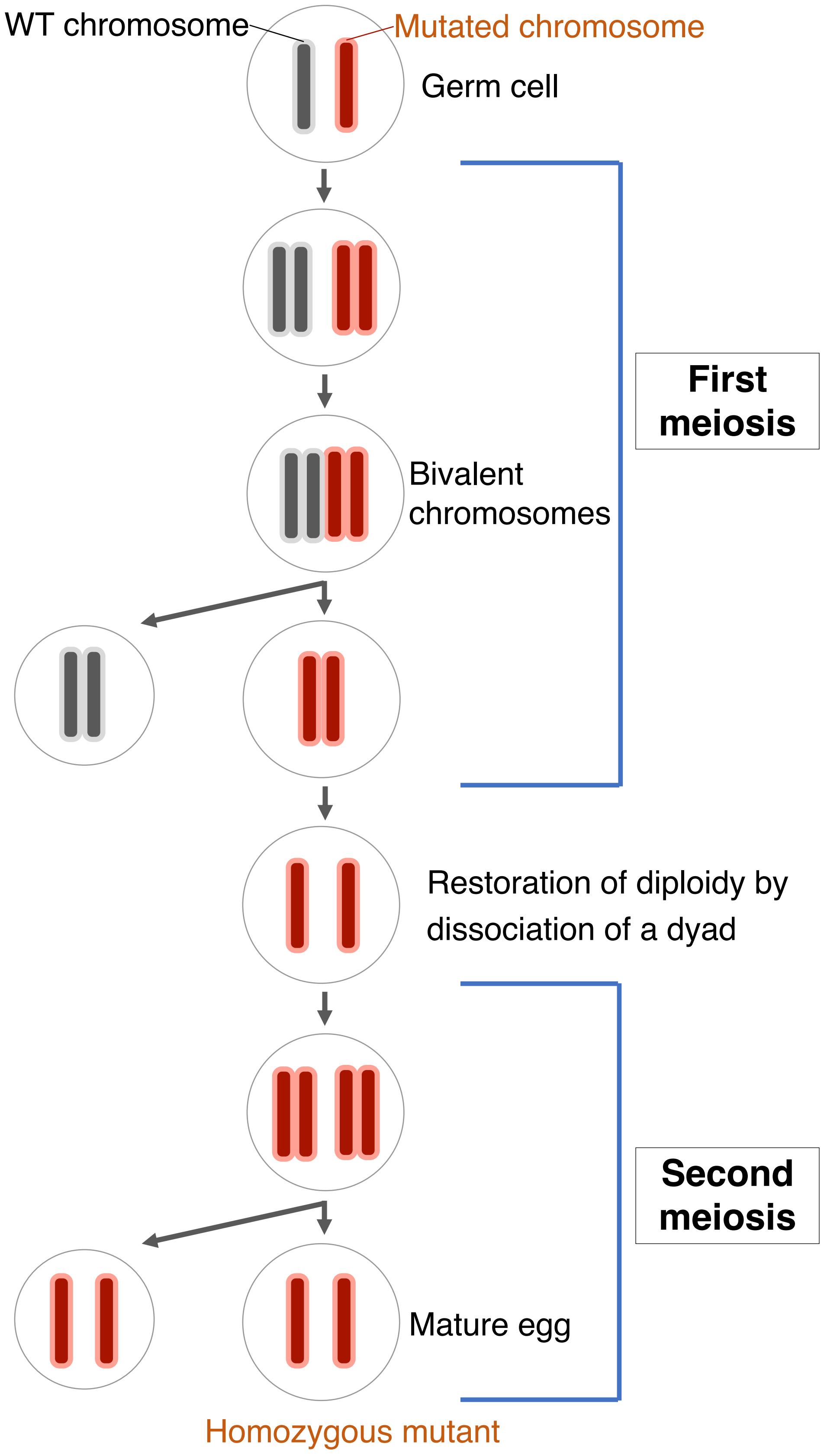


**Supplementary Fig. S4**

Proposed model for cytological process accounting for dominant generation of homozygous mutant in parthenogenetic tardigrades

**Supplementary Table S1.** Concentrations of Cas9 and glycerol used in the related studies.

| Cas9 conc. (μg/μL) | Glycerol conc. (%) | Notes |
|---|---|---|
| **4.0** | 20 | Maximum available Cas9 conc. |
| **3.3** | 16.5 | Used in the original DIPA-CRISPR for insects (ref. 25) |
| **3.0** | 15 | Used in this study. |
| **0.41** | 2.1 | Used in the CRISPR study for tardigrade somatic cells (ref. 24) |

**Supplementary Table S2.** Effects of glycerol concentration on the survival of the injected tardigrades.

| Glycerol conc. in injection solution (%) | # injected animals | # survived animals[a] (ratio) |
|---|---|---|
| **20** | 10 | 2 (20%) |
| **15** | 11 | 5 (45.5%) |
| **10** | 11 | 2 (18.2%) |
| **0** | 18 | 17 (94.4%) |

[a] examined at 24 h after injection

**Supplementary Table S3.** Hatchability of G1 eggs laid by G0 individuals carrying edited *tps-tpp*.

| G0 individuals ID (carrying mutation) | # G1 eggs laid by each G0 | # hatched G1 eggs (ratio) |
|---|---|---|
| **1 (1nt-ins)** | 8 | 0 (0%) |
| **2 (1nt-ins)** | 6 | 0 (0%) |
| **3 (1nt-ins)** | 2 | 0 (0%) |
| 4 **(1nt-ins)** | 3 | 0 (0%) |
| **5 (8nt + 3nt-del)** | 3 | 0 (0%) |
| **6 (484nt-del)** | 2 | 0 (0%) |
| **Total** | **24** | **0 (0%)** |

**Supplementary Table S4**. Hatchability of G1 eggs laid by G0 individuals without editing in *tps-tpp*.

| G0 individuals ID carrying no mutation | # G1 eggs laid by each G0 | # hatched G1 eggs (ratio) |
|---|---|---|
| 1 | 5 | 5 (100%) |
| 2 | 2 | 2 (100%) |
| 3 | 4 | 4 (100%) |
| 4 | 4 | 4 (100%) |
| 5 | 4 | 4 (100%) |
| 6 | 4 | 4 (100%) |
| 7 | 2 | 2 (100%) |
| 8 | 2 | 2 (100%) |
| 9 | 2 | 2 (100%) |
| 10 | 4 | 4 (100%) |
| 11 | 4 | 3 (75%) |
| 12 | 3 | 1 (33.3%) |
| 13 | 5 | 5 (100%) |
| 14 | 4 | 4 (100%) |
| 15 | 4 | 2 (50%) |
| 16 | 3 | 2 (66.7%) |
| 17 | 1 | 1 (100%) |
| **Total** | **57** | **51 (89.9%)** |

### Supplementary Table S5

The sequences of genome PCR primers (A), crRNAs (B) and ssODN (C).

#### **(A)** PCR primers

| Target genes | primers | sequences | notes |
|---|---|---|---|
| *white* (RvY_01244) | Forward | CTGGGAAATTATATGCTGATGGAGGG | Knock-out experiments |
| | Reverse | TAGCAGCAAGTAGAGGAGGACATGC | |
| *tps-tpp* (RvY_13060) | Forward | AGGAAGAGGAAGCTTCTGTGACGGA | |
| | Reverse | AGTCAATACTGATGGGGAAGGCGTC | |
| *white* (RvY_01244) | Forward | AGGCTCCATCTGTCAAACCAACCAG | Knock-in experiments |
| | Reverse | GATGTATTCCACGTCGAAGTCCGA | |

#### **(B)** crRNAs

| Target genes | crRNAs | sequences | notes |
|---|---|---|---|
| *white* (RvY_01244) | crRNA1 | CUGUUGGCGUCCCACAAUAC | Knock-out experiments |
| | crRNA2 | CCGAUACUCUGUCAAAGAGC | |
| | crRNA3 | GGAUUCAAGACCAUCCUGAA | |
| *tps-tpp* (RvY_13060) | crRNA1 | GCUUUUCCAACGAGACCUUG | |
| | crRNA2 | AGUGUGGAACCCAAUGACGU | |
| *white* (RvY_01244) | crRNA | GGUAAGAUCUGACUUUCCAG | Knock-in experiments |

#### **(C)** ssODN

| Target gene | ssODN sequence |
|---|---|
| *white* (RvY_01244) | CAAGTCCTTCCTGGATGAGATGGGCATCACCAACACGCACTGGTGGCTGGACTTCCTGGTTCTCTGCATGTTCTTTGTGGTTCTACGAGTCATCGCTTATCTAGTACTCCGCTGGAAAGTCAGATCTTACCGTTGAAGTCTCTTCTTCCTCCTATTTTCGTCTTGTTACTTCCCTGTTCAAAGGTAGCA |
